# Complementation can maintain a quasispecies of drug sensitive and resistant HIV

**DOI:** 10.1101/2020.04.20.051854

**Authors:** Laurelle Jackson, Sandile Cele, Gila Lustig, Jennifer Giandhari, Tulio de Oliveira, Richard A. Neher, Alex Sigal

## Abstract

HIV exists as multiple genotypes in a single infected individual referred to as a quasispecies. Here we reproduced a quasispecies by moderate selective pressure using an HIV reverse transcriptase inhibitor. The drug resistant genotype never completely supplanted the drug sensitive genotype, which stabilized at about 20 percent of viral sequences. Single-cell sequencing showed that resistant genotype frequency plateaued when cells were co-infected with sensitive and resistant genotypes, suggesting a sharing of viral proteins in co-infected cells (complementation) which masks genotypic differences. To test if complementation can confer phenotypic drug resistance, we co-transfected fluorescently labelled molecular clones of sensitive and resistant HIV and observed that genotypically sensitive virus from co-transfected cells was drug resistant. Resistant virus preferentially infected cells in tandem with drug sensitive HIV, explaining how co-infections of sensitive and resistant genotypes were initiated. Modelling this effect, we observed that a stable quasispecies could form at the experimental multiplicities of infection for the drug resistant and drug sensitive virus, showing that complementation can lead to a quasispecies in an HIV evolution experiment.

## Introduction

The HIV quasispecies consists of multiple viral genomes sampled from a compartment such as blood. One consequence of a quasispecies is diversity in the HIV Env gene. This allows HIV to escape neutralizing antibodies ***Frost et al.*** (***2005***); ***Rong et al.*** (***2009***). In the face of antiretroviral therapy, a quasispecies is formed which results in a mixture of HIV genomes that are resistant to different degrees to antiretroviral drugs (ARVs) ***Brenner et al.*** (***2002***); ***Allers et al.*** (***2007***); ***Samuel*** et al. (***2016***). A quasispecies may also allow HIV to maintain sufficient heterogeneity to escape effective T cell suppression ***Phillips et al.*** (***1991***); ***Goulder et al.*** (***2001***).

Due to the errors in reverse transcription, HIV replication generates a quasispecies where variants are at low frequencies around the main viral sequence ***Brenner et al.*** (***2002***); ***Coffin (1992***); ***Katz and Skalka*** (***1990***). Selection should act to clear less fit variants ***Rosenbloom et al.*** (***2012***), so how a stable quasispecies with different variants at high frequencies is maintained is unclear. One possibility is that different anatomical compartments apply different selective pressures, therefore leading to a diverse viral pool ***Paranjpe et al.*** (***2002***); ***Schnell et al.*** (***2010***). A mechanism to create a quasispecies that does not rely on the assumption of different compartments is complementation or phenotypic mixing ***Domingo et al.*** (***2012***); ***Hill et al.*** (***2012***). With this mechanism, co-infection and viral protein expression from two different viral genotypes in the same cell results in sharing of viral components. This mechanism is distinct from recombination, where two viral genomes co-packaged in the same cell recombine to form a new genome ***Levy et al.*** (***2004***). In complementation, a virus of genotype 1 may contain proteins from virus of genotype 2 and vice versa (Figure 1). If one of the genotypes has a fitness cost relative to the other, the difference in fitness will be masked ***Andino and Domingo*** (***2015***). This process has been postulated to operate in viruses ***Froissart et al.*** (***2004***); ***Vignuzzi et al.*** (***2006***) including HIV ***Mo et al.*** (***2004***); ***Gelderblom et al.*** (***2008***). There is no known mechanism which prevents one HIV genotype packaging viral proteins such as reverse transcriptase (RT) from another HIV genotype if both genotypes are expressed in the same cell, since RT molecules mix in the cell cytoplasm ***Freed*** (***2001***).

**Figure 1.**
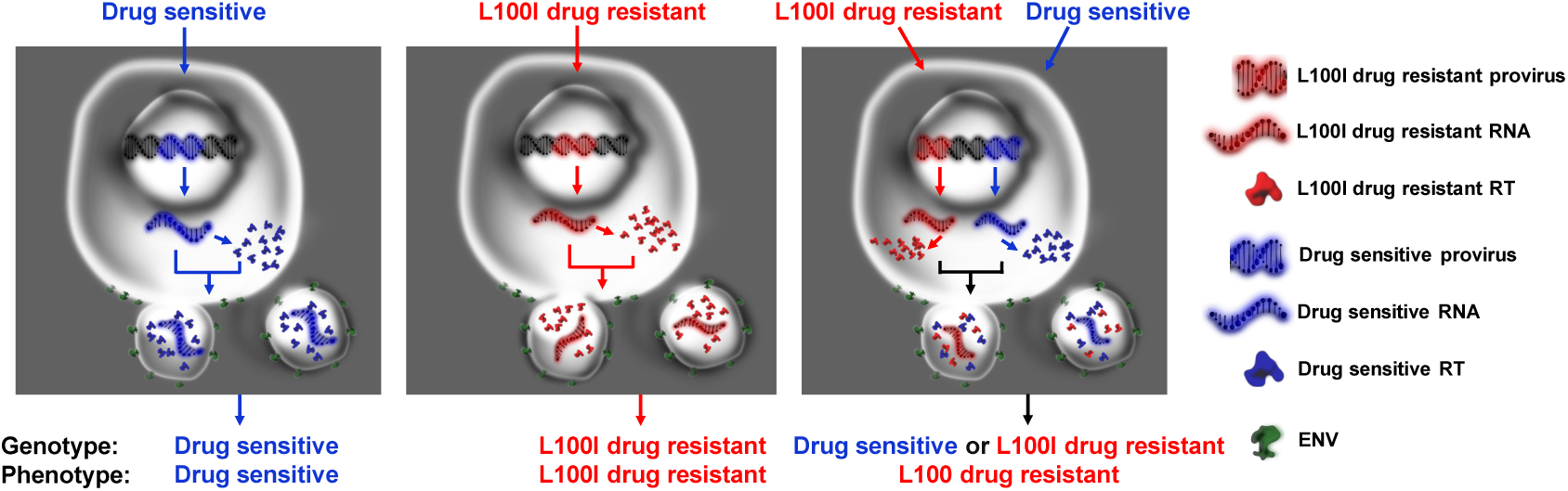
Schematic of how drug sensitive and drug resistant HIV could co-exist by complementation in the face of selective pressure applied by a reverse transcriptase inhibitor. When drug sensitive and drug resistant virus infect separate cells, the virus genotype corresponds to the virus phenotype (left two panels). When a drug sensitive and drug resistant virus infect the same cell, the virions produced will have either a drug sensitive or drug resistant genotype but similar numbers of drug resistant and drug sensitive reverse transcriptase molecules, and hence the same phenotypic drug resistance (right panel).

Here we used *in vitro* HIV evolution in the presence of the RT inhibitor efavirenz (EFV) to determine whether a quasispecies can be formed in a homogeneous infection environment. We used a concentration of drug which allowed drug sensitive virus to replicate, but conferred a strong fitness advantage to drug resistant virus. We observed that drug resistance evolved during infection in the face of EFV. Initially, viral genotypes with drug resistance mutations expanded rapidly. However, once the fraction of drug resistance mutant infected cells reached approximately 80%, the frequency of drug sensitive virus stabilized. This correlated with co-infection of drug sensitive and drug resistant genomes, and such co-infection led to virions with drug sensitive and drug resistant genotypes to display a similar level of EFV resistance. Therefore, complementation can maintain a quasispecies of viral variants having different fitness in the presence of drug.

## Results

To test whether a quasispecies can be reproduced *in vitro*, we infected cells from an HIV reporter cell line in the face of drug pressure from EFV. As the reporter cells we used the the RevCEM E7 clone (***Boullé et al.*** (***2016***); ***Jackson et al.*** (***2018***)) from the RevCEM GFP reporter cell line. These cells express GFP in the presence of the HIV Rev protein (***Wu et al.*** (***2007***)), with the maximum frequency of GFP positive cells being about 70 percent in the E7 clone (***Jackson et al.*** (***2018***)). Infection was initiated with 4 days (approximately 2 viral cycles) of infection in the absence of drug. Drug was then added (day 0) and infection measured every two days. After the frequency of infected cells was recorded, infection was diluted to 2% of the total cell population by addition of uninfected cells. This enabled us determine the degree to which infection could expand in each 2 day infection cycle without exhausting the population of uninfected target cells.

The EFV concentration used was 20nM. This allowed drug sensitive (wildtype) HIV to persist (Figure 2A). The number of cells infected in a two day infection cycle starting at 2% infected cells began to increase at day 6 post-addition of EFV selective pressure (Figure 2A). An increase in the number of infected cells is expected with the evolution of drug resistance.

**Figure 2.**
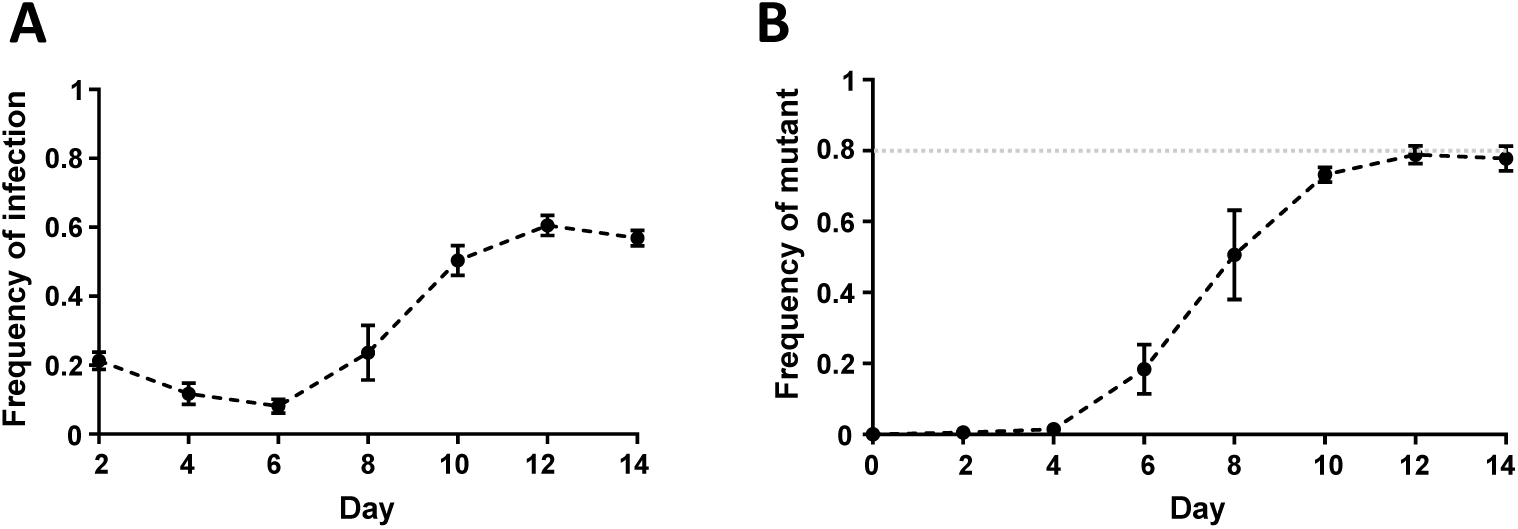
EFV resistant mutants do not completely supplant drug sensitive viral sequences in the face of EFV selection. (A) Frequency of infection as a function of time in the presence of 20nM EFV. Cells were cultured for two infections cycles before the addition of drug (day 0). Thereafter, the frequency of infection was measured every 2 days. Mean ± SEM of 3 independent experiments. (B) Frequency of EFV resistant mutants as a function of time derived from illumina sequencing of the cell populations from (A). Mean ± SEM of 3 independent experiments. Dotted line marks frequency of 0.8.

To examine whether evolution of drug resistance did take place, we sequenced the infected cell population and determined the fraction of sequences with drug resistance mutations. Sequencing of the HIV DNA from the infected cell population showed that starting day 6 post-drug addition, mutations in the HIV RT gene which confer resistance to EFV were detected (Figure 2B, see Figure S1 for individual independent experiments and the mutations which arose). Predominant among these was the L100I mutation (Figure S1), conferring moderate resistance to EFV ***Jackson et al.*** (***2018***). The frequency of mutant genotypes reached about 80% on day 10. The combined frequency of the mutants stabilized at about 80% on day 12 and day 14. Drug sensitive HIV accounted for the remaining 20 percent of sequences in the population.

To examine whether the plateau in the frequency of drug resistant mutants was correlated with co-infection of the same cell with wildtype and mutant HIV, we single-cell sequenced HIV DNA in 30 to 60 cells by sorting GFP positive infected cells into wells of a multi-well plate at 1 cell per well, then lysing and amplifying the RT region followed by illumina sequencing (Figure 3). At day 0, before selective pressure was applied, all infected cells showed wildtype HIV genomes. At day 6, 35% of cells had EFV resistance mutations. On day 8 and day 14, where the mutant frequency stabilized at about 80% in the population measurement, the majority of cells were infected by EFV resistant mutants. On all days where drug resistant mutant infection was present, individual cells also contained wildtype sequences. At later time points, there was an increase in cells infected with multiple drug resistance mutant genomes. There was greater than expected frequency of the G190A mutation relative to the population data, but this was mostly explained by variation in mutant frequencies between experiments (Figure S1), with the single-cell sequences originating in an experiment which showed higher frequencies of G190A at the population level (Figure S1, experiment 3).

**Figure 3.**
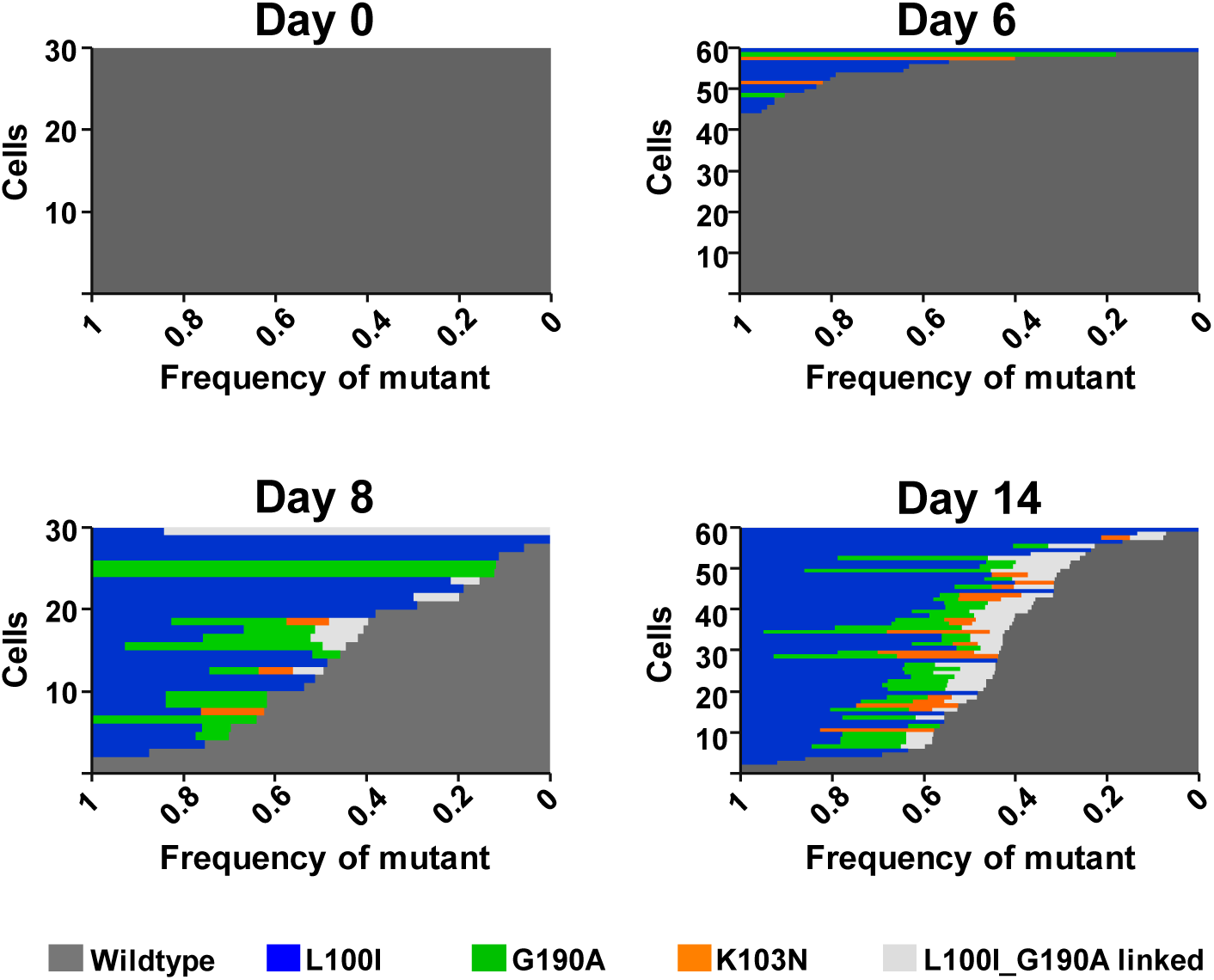
Sequencing of viral genomes in single cells shows increasing frequency of co-infected cells with time. Shown are the relative frequencies of wildtype and mutant HIV genomes per cell for the day 0, day 6, day 8 and day 14 post-EFV addition. Frequency is on the x-axis, cell number on y-axis. Cells were ranked by lowest to highest wildtype frequency. Dark grey is the wildtype genotype, blue is the L100I mutant, orange is the K103N mutant, green is the G190A mutant, and light grey is the L100I/G190A linked double mutant.

To directly test whether an EFV resistant mutant HIV can complement wildtype, we used transfection of molecular viral clones consisting of plasmids expressing a mutant and wildtype virus in conjunction with a fluorescent protein. Upon transfection, molecular clones produce fully functional replicating virus which can be tested for resistance to EFV. This system offers direct control over co-expression of viral proteins in the same cell, as wildtype and mutant virus expressing plasmids can be transfected separately or mixed for efficient co-transfection. We transfected molecular clones expressing either wildtype CFP labeled virus (***Levy et al.*** (***2004***)) or YFP labelled virus in which we replaced the RT region with the L100I mutant into a virus producer cell line. The producer cells showed either CFP or YFP fluorescence, depending on the transfected virus (Figure 4A, left panel). When we co-transfected the molecular clones, we observed dual CFP and YFP expression from most fluorescent cells, indicating both viral strains being expressed from the same cell (Figure 4A, right panel). We collected viral supernatant from each of the three conditions, filtered the viral supernatant to avoid any carryover of transfected cells, and used it to infect the lymphocytic MT4 cell line (see Figure S2 for gating strategy). To quantify infection under equivalent detection conditions, we mixed the supernatants from the single genotype transfected, wildtype only CFP expressing cells, and single genotype transfected, mutant only YFP expressing cells. We compared this infection to that of supernatant from the wildtype/mutant co-transfected cells. The EFV sensitivity of each genotype in a mixed infection could then be tracked based on the decrease of CFP (wildtype) expressing cells or YFP (mutant) expressing cells with escalating EFV concentration (see Figure S2 for gating strategy). Cells infected with virus made from mixed single genotype transfections were either EFV sensitive (wildtype, CFP expressing cells) or resistant (mutant, YFP expressing cells) (Figure 4B, left panel). In contrast, virus from the co-transfection showed EFV resistance for both the wildtype CFP expressing genotype and the YFP expressing mutant genotype (Figure 4B, right panel). For wildtype, resistance gained was comparable to the YFP resistant mutant. This indicates that the wildtype virus was able to complement with the mutant virus.

**Figure 4.**
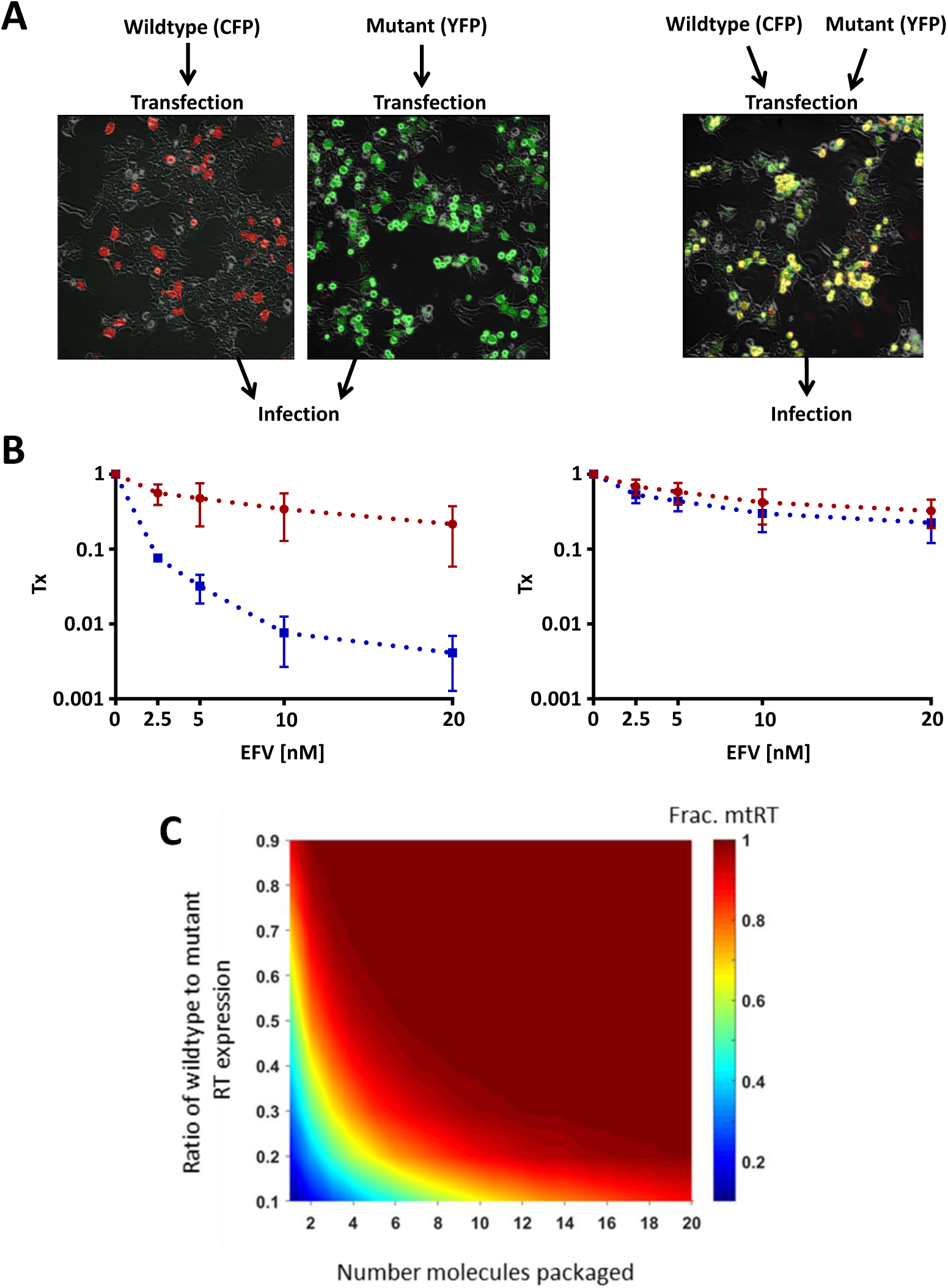
Co-transfection of wildtype and mutant molecular clones yields resistant virus independent of genotype. (A) CFP expressing wildtype and YFP expressing L100I EFV resistant mutant molecular clones were either transfected separately (left) or co-transfected (right) into a virus producer cell line. Shown are images after producer line transfection, with CFP in red and YFP in green. Co-transfected cells which express both fluorescent proteins are yellow. (B) Sensitivity of cell-free virus collected from the transfections to EFV. Sensitivity was measured as the transmission index (Tx), the ratio of the number of infected cells in the presence of EFV divided by the number of infected cells in the absence of EFV for each genotype. Wildtype (CFP labelled) and mutant (YFP labelled) genotypes were determined by the corresponding fluorescence. Left panel shows a mix of wildtype and mutant virus from separate transfections, while right panel shows virus from co-transfection. Red and blue points are the Tx values of mutant and wildtype genotypes respectively (mean ± std of 3 independent experiments). (C) Predicted fraction of virions containing at least one mutant RT molecule as a function of the ratio of genomic copies of drug sensitive versus resistant virus and the number of RT molecules packaged.

We calculated the probability that at least one molecule of mutant RT is packaged per virion in co-infected cells as a function of the number of RT molecules packaged per virion and the ratio of wildtype to mutant viral genomes, assuming each genome produces the same number of RT proteins. At low numbers of RT molecules and high number of wildtype genomes, the probability that mutant RT will be packaged is low and complementation should be rare. At the reported numbers of RT molecules per virion (roughly 50) (***Panet and Kra-Oz*** (***1978***); ***Bauer and Temin*** (***1980***)), the fraction of co-packaging virions and therefore complementation is high (Figure 4C).

Whether co-infection is common has been an area of active debate (***Jung et al.*** (***2002***); ***Law et al.*** (***2016***); ***Josefsson et al.*** (***2011***, 2013)). However, it was previously observed that HIV preferentially co-infects cells at relatively low infection frequencies (***Dang et al.*** (***2004***); ***Del Portillo et al.*** (***2011***)), possibly due to heterogeneity in the cell population, cooperativity between viruses, or other factors. To test for this, we infected a RevCEM clone selected for lack of GFP expression (Materials and methods) with CFP expressing wildtype virus or YFP expressing L100I mutant (Figure 5A). We observed co-infection frequencies which were approximately one-order of magnitude higher than predicted under the assumption that infections were independent (Figure 5B, see Figure S3 for flow cytometry plots). This effect was even stronger in peripheral blood mononuclear cells (Figure S4) when these cells were infected in the presence of a cell line expressing the DC-SIGN receptor. DC-SIGN binds HIV on the surface. Infection with surface bound virus is efficient (***Kim et al.*** (***2018***)) and is important in environments such as the germinal center of the lymph node (***Fletcher et al.*** (***2014***)). Hence, a high frequency of co-infection need not be common overall and may be localized to specific environments.

**Figure 5.**
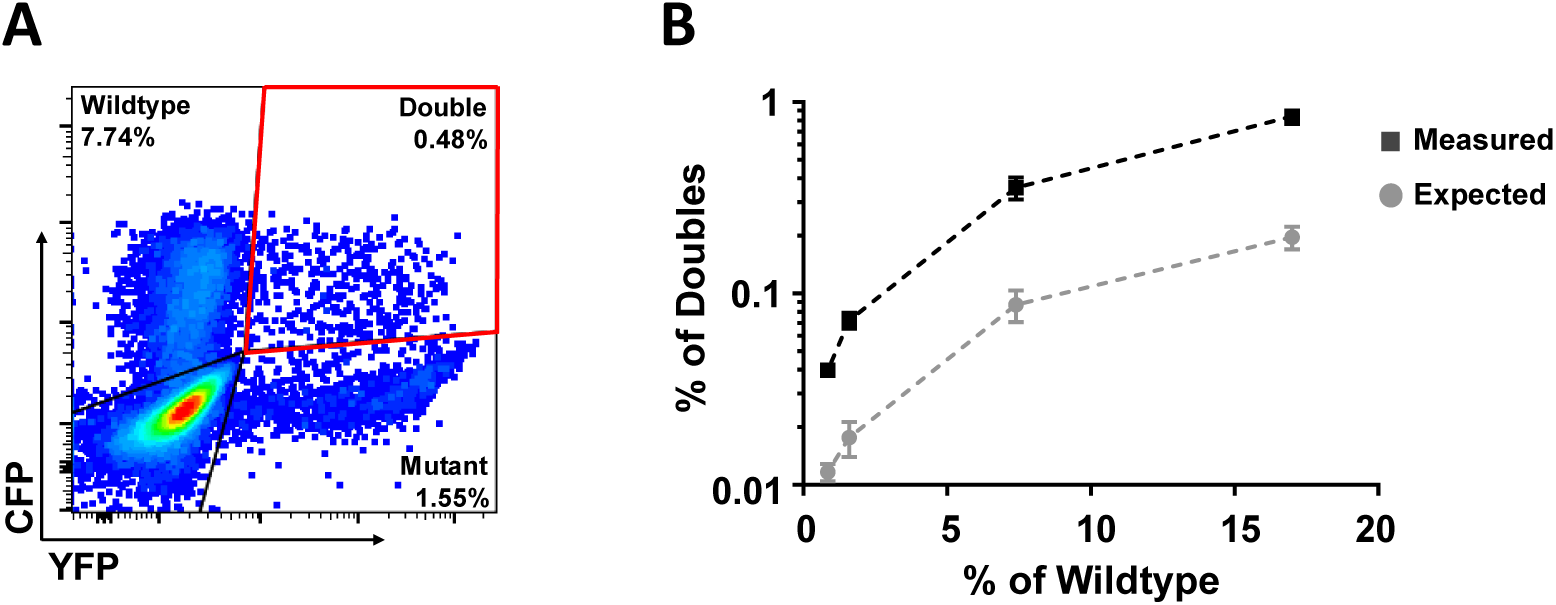
Double infection is more frequent than expected assuming independence. (A) CFP labelled wildtype HIV the YFP labelled mutant were used to infect non-reporter C7 target cells in the absence of drug. Infection was titrated to yield different wildtype frequencies, and the percent of double infected cells detected in the top right quadrant. (B) The measured (black squares) versus expected (grey squares) percentage of double infection at different wildtype frequencies. Expected probability of double infection was calculated as the product of the wildtype and mutant single infection frequencies. Mean ± SEM from 3 independent experiments.

To investigate whether a stable quasispecies is predicted to occur given experimentally measured parameter values, we simulated the frequencies of wildtype, mutant and co-infected cells through time (Materials and methods). The experimentally measured values for the multiplicity of infection were calculated as ***R***_0_***I***_*input*_/***T***, where ***R***_0_ is the basic reproductive ratio for wildtype or mutant, measured at an input of 0.2% infected cells (Figure S5, Tables S1 and S2), ***I***_*input*_ the input number of wildtype or mutant infected cells, and ***T*** the number of uninfected cells). The assumption was made in the model that a cell co-infected with mutant and wildtype virus would produce half of the virions with a wildtype genotype and half with mutant, and that these would contain sufficient drug resistant RT to be phenotypically mutant, consistent with a high number of RT molecules per virion (Figure 4C). We observed that at the experimentally measured values, cells infected with mutant virus did not entirely supplant wildtype infected cells. There was a high and stable frequency of both co-infected cells and a lower but stable frequency of wildtype only infected cells (Figure 6A). Examining these effects as a function of multiplicity of infection of the mutant virus showed that the frequency of wildtype infected cells (sum of co-infections with mutant and wildtype only) were absent at mutant multiplicities of infection below approximately 2 and sharply increased thereafter. This effect can be explained by the requirement of wildtype infected cells to be co-infected by mutant at high frequencies for the wildtype not to be outcompeted.

**Figure 6.**
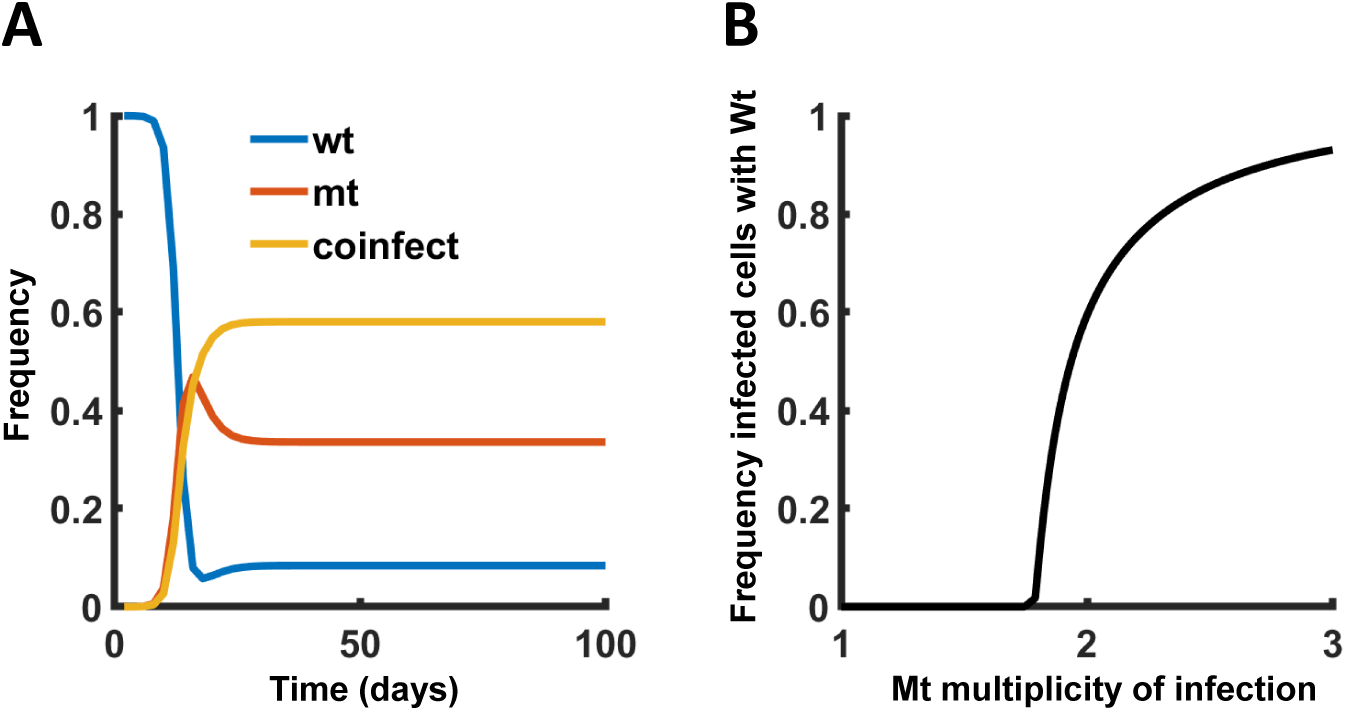
Model of complementation shows that a stable quasispecies is predicted at the experimentally measured parameter values. (A) Predicted frequencies of wildtype infected (wt, blue line), drug resistant mutant infected (mt, red line), and coinfected wildtype-mutant (coinfect, yellow line) cells over 50 viral cycles (approximately 100 days). (B) The fraction of cells infected with wildtype after 500 viral cycles, both as wildtype only and co-infected with mutant, as a function of mutant multiplicity of infection. Multiplicity of infection is calculated as 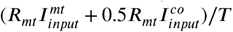, where ***R***_*mt*_ is the measured ***R***_0_ for mutant, 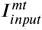 is the input number of mutant infected cells, 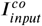 is the number of input co-infected cells, and ***T*** is the number of uninfected target cells.

## Discussion

We observed that a quasispecies is formed in the face of EFV selective pressure, where wildtype HIV persists despite a fitness disadvantage relative to drug resistant virus. The necessary condition for complementation to occur *in vivo* is expression of multiple viral genotypes in the same cell. Multiple infections can occur by cell-to-cell spread, a directed mode of HIV transmission efficiently delivering HIV from the donor to target cell (***Jolly et al.*** (***2004***); ***Sattentau*** (***2008***); ***Sigal et al.*** (***2011***); ***Jackson*** et al. (***2018***); ***Agosto et al.*** (***2015***); ***Abela et al.*** (***2012***); ***Baxter et al.*** (***2014***); ***Boullé et al.*** (***2016***); ***Hübner*** et al. (***2009***); ***Del Portillo et al.*** (***2011***); ***Iwami et al.*** (***2015***); ***Duncan et al.*** (***2013***); ***Groot et al.*** (***2008***); ***Sherer et al.*** (***2007***); ***Zhong et al.*** (***2013***)), likely to to occur in localized environments with minimal flow to disrupt cellular interactions (***Sourisseau et al.*** (***2007***)). HIV can be transmitted by cell-to-cell spread to multiple cells at once (***Rudnicka et al.*** (***2009***)). Therefore, one cell can be infected by multiple HIV genomes if there are multiple HIV transmitting cells in its vicinity.

The frequency of multiple HIV proviruses per cell has been controversial, with some studies showing multiple infection per cell (***Jung et al.*** (***2002***); ***Law et al.*** (***2016***); ***Gratton et al.*** (***2000***)). Other studies did not show multiple infections at a frequency greater than that predicted by the Poisson distribution (***Josefsson et al.*** (***2011***, 2013)). Given that many proviruses are not expressed (***Bruner*** et al. (***2016***)) and that cells with multiple infections may be more likely to express HIV and be infectious (***Wodarz and Levy*** (***2017***)), it is possible that the multiplicity of infection may be higher than predicted by Poisson in cells where HIV is actively replicating (***Pardons et al.*** (***2019***)). Other possibilities where localized multiple infection per cell can occur is in cell subsets (***Banga et al.*** (***2016***)) where HIV infection is particularly efficient in lymph nodes or gut (***Fletcher et al.*** (***2014***); ***Brenchley et al.*** (***2004***); ***Deleage et al.*** (***2016***)).

Complementation can reduce the fitness of the viral population by preventing the selection of more fit genotypes (***Froissart et al.*** (***2004***)). There is evidence that the quasispecies stabilized by complementation is beneficial for the fitness of the population, since this allows deleterious variants to share components and result in a more functional virion, or keep a heterogeneous pool of virus which can react to rapid changes in the infection environment (***Vignuzzi et al.*** (***2006***); ***Domingo et al.*** (***2012***); ***Lauring et al.*** (***2013***); ***Andino and Domingo*** (***2015***)). Interestingly, the majority mutant HIV in this study was coinfected with wildtype. This may indicate that wildtype infection may make the cells more permissive for the L100I mutant.

Complementation may be one example of how the quantitative parameters of infection, and specifically the infecting dose (***Moyano et al.*** (***2020***); ***Sigal et al.*** (***2011***); ***Wodarz and Levy*** (***2017***)), may change the nature of infection and how it responds to inhibitors. The relatively high multiplicity of infection required for complementation need not be pervasive nor occur due to a very high ***R***_0_, but may also be the result of a high concentration of infected donor relative to yet uninfected target cells. This study shows that HIV complementation can occur and lead to a quasispecies under conditions where the multiplicity of infection can be experimentally controlled. The extent this effect occurs *in vivo* is yet to be determined.

## Methods and Materials

### Inhibitors, viruses and cell lines

The antiretroviral EFV, Raji cells and Raji-DC cells were obtained from the AIDS Research and Reference Reagent Program, National Institute of Allergy and Infectious Diseases, National Institutes of Health. RevCEM cells from Y. Wu and J. Marsh; MT-4 cells from D. Richman and HIV molecular clone pNL4-3 from M. Martin. The NL4-3YFP and NL4-3CFP molecular clones were gifts from D. Levy. Cell-free viruses were produced by transfection of HEK293 cells with pNL4-3 using TransIT-LT1 (Mirus) or Fugene HD (Roche) transfection reagents. Virus containing supernatant was harvested after two days of incubation and filtered through a 0.45/*mu*m filter (Corning). The number of virus genomes in viral stocks was determined using the RealTime HIV-1 viral load test (Abbott Diagnostics). The L100I mutant was evolved by serial passages of wildtype NL4-3 in RevCEM cells in the presence of 20nM EFV. After 18 days of selection, the RT gene was cloned from the proviral DNA and the mutant RT gene was inserted into the NL4-3 molecular clone. RevCEM clone E7 and G2 used in this study were generated as previously described ***Boullé et al.*** (***2016***). Briefly, the E7 clone was generated by subcloning RevCEM cells at single cell density. Surviving clones were subdivided into replicate plates. One of the plates was screened for the fraction of GFP expressing cells upon HIV infection using microscopy, and the clone with the highest fraction of GFP positive cells was selected. As with the E7 clone, for the generation of the RevCEM clone C7, cells were split into single cell density and one of the plates were screened for the lowest fraction of GFP expression upon HIV infection. The clone with lowest fraction of GFP was selected. All cell lines not authenticated, and mycoplasma negative. Cell culture and experiments were performed in complete RPMI 1640 medium supplemented with L-Glutamine, sodium pyruvate, HEPES, non-essential amino acids (Lonza), and 10 percent heat-inactivated FBS (Hyclone). For transfections of CFP and YFP viruses, NL4-3YFP and NL4-3CFP were added to Fugene HD (Roche) for single transfections co-transfection into HEK293 cells.

### Primary cells

Blood for PBMC was obtained from HIV negative blood donors with no TB symptoms. Informed consent was obtained from each participant, and the study protocol approved by the University of KwaZulu-Natal Institutional Review Board (approval BE083/18). PBMCs were isolated by density gradient centrifugation using Histopaque 1077 (Sigma-Aldrich) and cultured at 10^6^ cells/ml in complete RPMI 1640 medium supplemented with L-Glutamine, sodium pyruvate, HEPES, non-essential amino acids (Lonza), 10 percent heat-inactivated FBS (GE Healthcare Bio-Sciences), and IL-2 at 5 ng/ml (PeproTech). Phytohemagglutinin at 10 /*mu*g/ml (Sigma-Aldrich) was added to activate cells.

### Staining and low cytometry

The frequency of RevCEM E7 GFP positive cells was detected on on a FACSCaliber (BD Biosciences) machine using the 488 laser lines. The number of CFP and YFP positive cells was determined on an FACAriaIII (BD Biosciences) using the 405nm and 488nm laser lines. Results were analysed with FlowJo 10.0.8 software. For single cell sorting to detect the number of HIV DNA copies per cell, GFP positive cells were single cell sorted using 85 micron nozzle in a FACSAriaIII machine into 96 well plates (Biorad) containing 30*µ*l lysis buffer (2.5*µ*l 0.1M Dithiothreitol, 5*µ*l 5 percent NP40 and 22.5*µ*l molecular biology grade water.

### Deep sequencing

For single cells, the cells were lysed and DNA was kept suspended in the lysis buffer. For populations of cells, genomic DNA was extracted using the QIAamp DNA mini kit (Qiagen). Phusion hot start II DNA polymerase (New England Biolabs) PCR reaction mix (10*µ*l 5X Phusion HF buffer, 1*µ*l dNTPs, 2.5*µ*l of the forward primer, 2.5*µ*l of the reverse primer, 0.5*µ*l Phusion hot start II DNA polymerase, 2.5*µ*l of DMSO and molecular biology grade water to 50*µ*l reaction volume) was added to the lysed single cells or extracted genomic DNA of cell populations. Two rounds of PCR were performed. The first-round reaction amplified a region of the RT gene in the proviral DNA using the forward primer 5’ tcgtcggcagcgtcagatgtgtataagagacagTTAATAAGAGAACTCAAGATTTC 3’ and reverse primer 5’ gtctcgtgggctcggagatgtgtataagagacagCCCCACCTCAACAGATGTTGTC 3’. Non-capitalized portion of the primers represent the Nextera® XT Index Kit adaptors. Cycling program was 98°C for 30 seconds, then 35 cycles of 98°C for 10 seconds, 50°C for 30 seconds and 72°C for 15 seconds with a final extension of 72°C for 5 minutes. 1*µ*l of the first round product was then transferred into a PCR mix as above, with second round Nextera XT Index Kit adaptor primers (forward 5’ tcgtcggcagcgtcagatgtgtataagagacag 3’, reverse 5’ gtctcgtgggctcggagatgtgtataagagacag 3’). The second round PCR amplified a 400bp product which was then visualized on a 1% agarose gel. The PCR amplicon was gel extracted using the QIAquick gel extraction kit (Qiagen). Illumina indices were attached to the amplicon with the Nextera XT Index Kit and deep sequenced using the Illumina Miseq. Fast-q files were analysed in Geneious. Both 5’ and 3’ ends were trimmed with an error probability limit of 0.1. Drug resistant mutations were found based on a minimum variant frequency of 0.01 for populations of cells and 0.05 for single cells. The maximum variant P-value was 10^−6^.

### Evolution

Experiments were initiated with a cell-free infection, where 10^6^ cells/ml RevCEM E7 were infected with 2 ×10^8^ NL4-3 viral copies/ml (roughly 20ng p24 equivalent) for 2 days. Infected cells from the cell-free infection were used as the donors and cocultured with 10^6^ cells/ml target cells. After two days of infection, 2% of the infected cells were added to uninfected target cells and cocultured for a 2-day cycle (day 0). Thereafter, 2% of resuspended infected cells were added to uninfected targets in the presence of EFV and co-cultured for each 2-day cycle.

### Frequency of CFP and YFP double infections

10^6^ RevCEM clone C7 cells/ml were infected with 2×10^8^ wildtype NL4-3CFP viral copies/ml (roughly 20ng p24 equivalent) for 2 days. Infected cells from the cell-free infection were used as the donors and cocultured with 10^6^ cells/ml C7 uninfected target cells at ratios of 1:5, 1:10, 1:50 and 1:100. The L100I mutant NL4-3YFP virus was added at 1:50 dilution (roughly 2ng p24 equivalent) to each wildtype NL4-3CFP donor to target condition. Cells were incubated for 2 days and analysed by flow cytometry. For the Raji and Raji-DC experiments, Raji cells and Raji-DC cells were added to PBMCs at ratios of 1:2 and then infected with 2×10^8^ NL43-YFP and NL43-CFP viral copies/ml (roughly 20ng p24 equivalent) for 2 days and analyzed by flow cytometry.

### Simulation of the fraction of cells containing mutant RT

A stochastic simulation was performed to find the probability of a virus packaging at least one mutant RT molecule as a function of the total number of RT molecules packaged by one virus and the fraction of RT molecules in the cell produced by the mutant RT gene. A vector of random numbers with number of entries equal to the number of total RT molecules packaged was chosen using Matlab from a uniform distribution between 0 and 1. If at least one of the numbers was less than 1-*F*_*wt*_, where *F*_*wt*_ is the wildtype frequency, the iteration was scored as containing at least one mutant RT. 10^4^ iterations were performed for each combination of total RT molecules and *F*_*wt*_, and the frequency of iterations with at least one mutant RT graphed.

### Simulation of wildtype, mutant and co-infected cell frequencies

Measured or set parameter values were:

- Total number of cells (***T***) = 6 × 10^6^
- Input of total infected cells at each viral cycle (*i*) = 1.2 × 10^5^
- Input of mutant 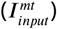, wildtype 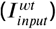, and co-infected 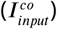 cells infected cells at viral cycle *i* are ***I***_*mt*_ = *F*_*mt*_***I, I***_*wt*_ = *F*_*mt*_***I, I***_*co*_ = *F*_*co*_***I*** where *F*_*mt*_, *F*_*wt*_, *F*_*co*_ are the fraction of mutant infected, wildtype infected, and co-infected cell and the end of infection cycle *i* − 1
- Mutant ***R***_0_ (***R***_*mt*_) = 162
- Wildtype ***R***_0_ (***R***_*wt*_) = 26

The multiplicity of infection for mutant and wildtype at infection cycle *i* with complementation was calculated as

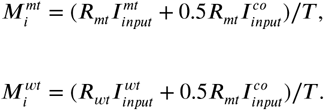

The probabilities of cells being infected with mutant and wildtype 10^−6^ were:

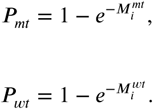

The probability of co-infection was therefore:

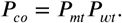

The number of co-infected cells in cycle *i* was:

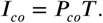

The number of mutant only and wildtype only infected cells in cycle *i* was:

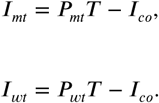

The fraction of cells in each infection state (mutant, wildtype, co-infected) was therefore ***I***_*mt*_/***T***, ***I***_*wt*_/***T***, ***I***_*co*_/***T*** respectively.

## Acknowledgments

This work was supported by Research Group Leader funding from the Max Planck Society (AS).

---

**Figure S1.**
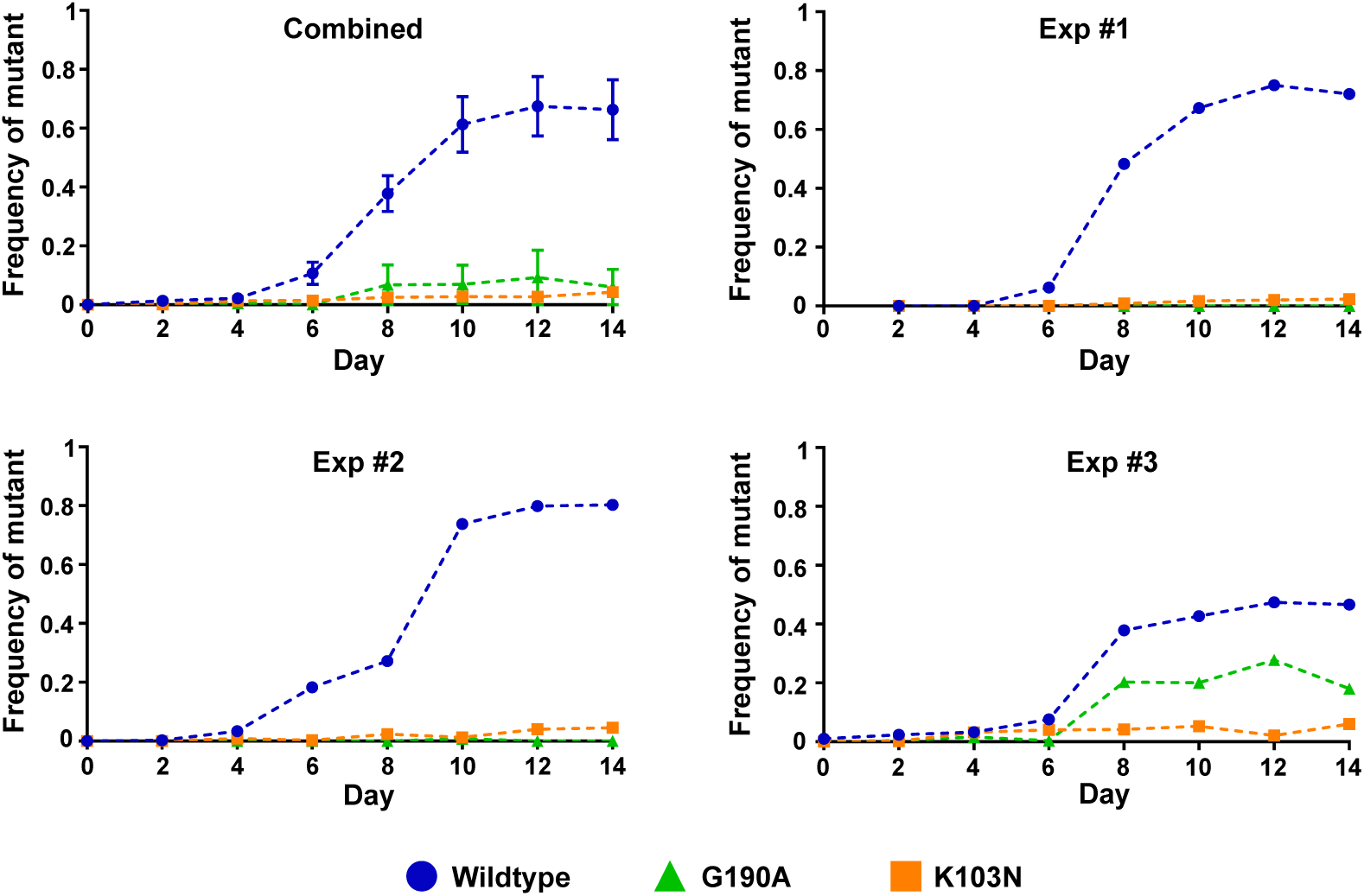
Frequency of individual EFV resistance mutants in the face of EFV selection in individual experiments. Combined graph is the mean ± SEM of individual experiments.

**Figure S2.**
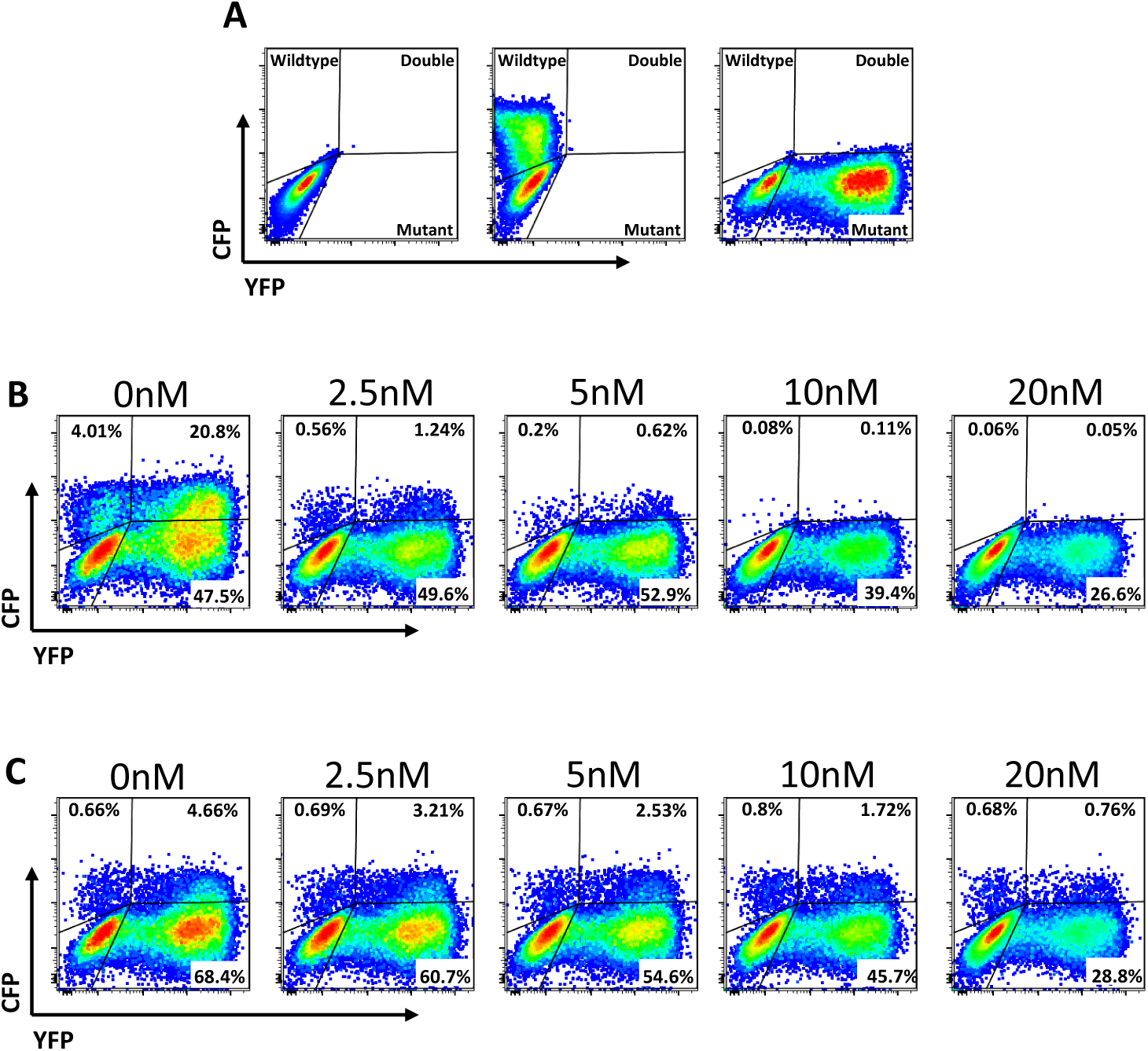
Gating strategy to detect EFV sensitivity of wild type and mutant HIV from singly transfected and co-transfected molecular clones. For single transfections, CFP labelled wildtype and YFP labelled mutant virus was mixed 1:1 before the infection. (A) Uninfected (left panel), CFP wildtype virus only (middle panel) or YFP mutant virus only (right panel) controls. (B) Infection with virus from single transfections mixed 1:1. X-axis is YFP (mutant), y-axis is CFP (wildtype) infected cells. EFV concentration used is shown above each plot. The total frequency of wildtype infected cells is the sum of the frequencies in the upper left and upper right quadrants, while the frequency of mutant infected cells is the sum of the lower right and upper right quadrants. (C) Infection of virus derived from the co-transfection. EFV concentration used is shown above each plot, and gating is as in (B).

**Figure S3.**
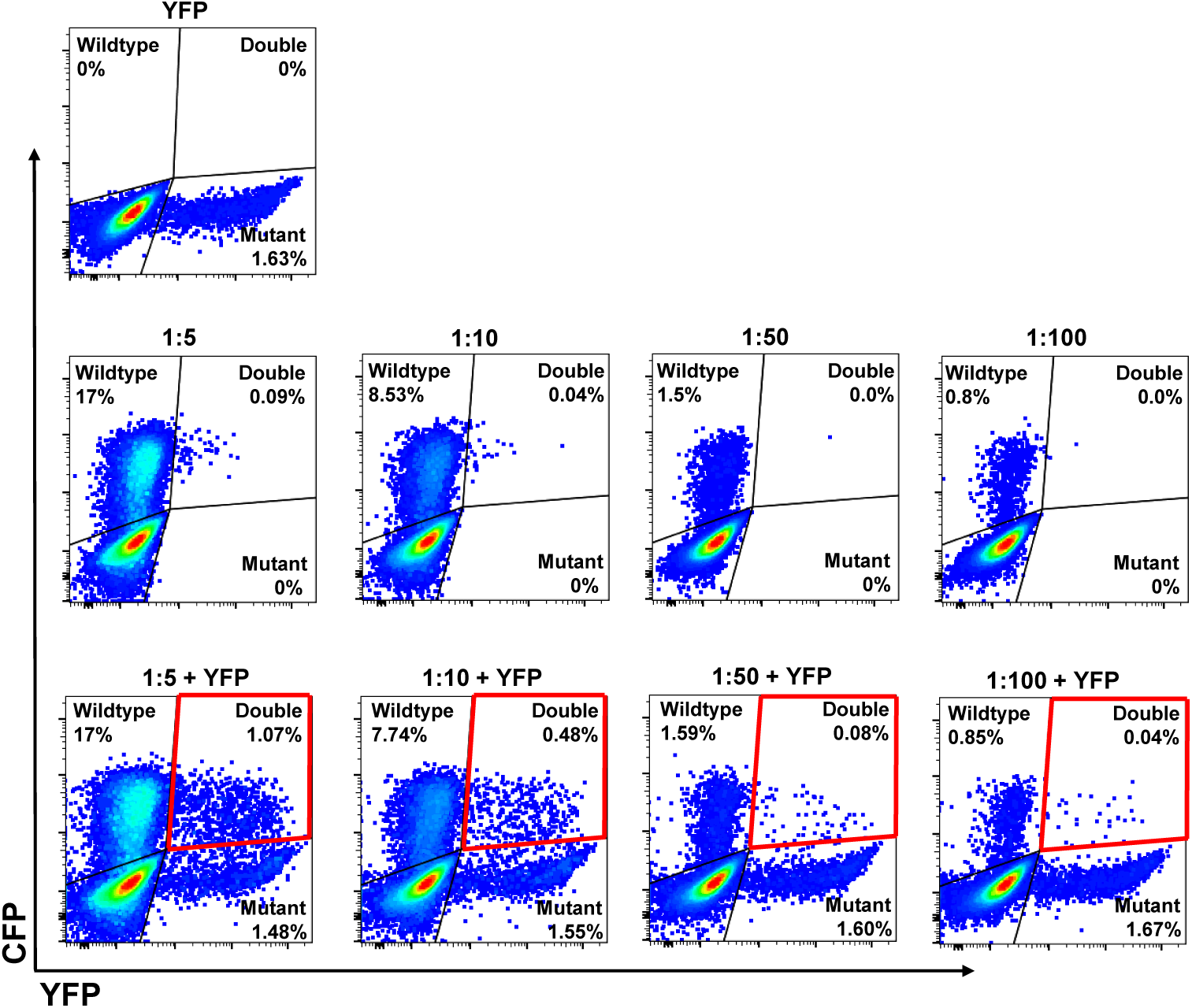
Multiple infection is more frequent than expected by chance: Flow cytometry data. Top row shows C7 cells infected with labelled YFP mutant viral supernatant. Middle row shows cells infected with CFP labelled wildtype using an input of infected cells at a dilution shown above each plot. Bottom row shows cells infected with CFP labeled wildtype using the same input of infected cells as the middle row, but with the addition of the same amount of labelled YFP mutant viral supernatant.

**Figure S4.**
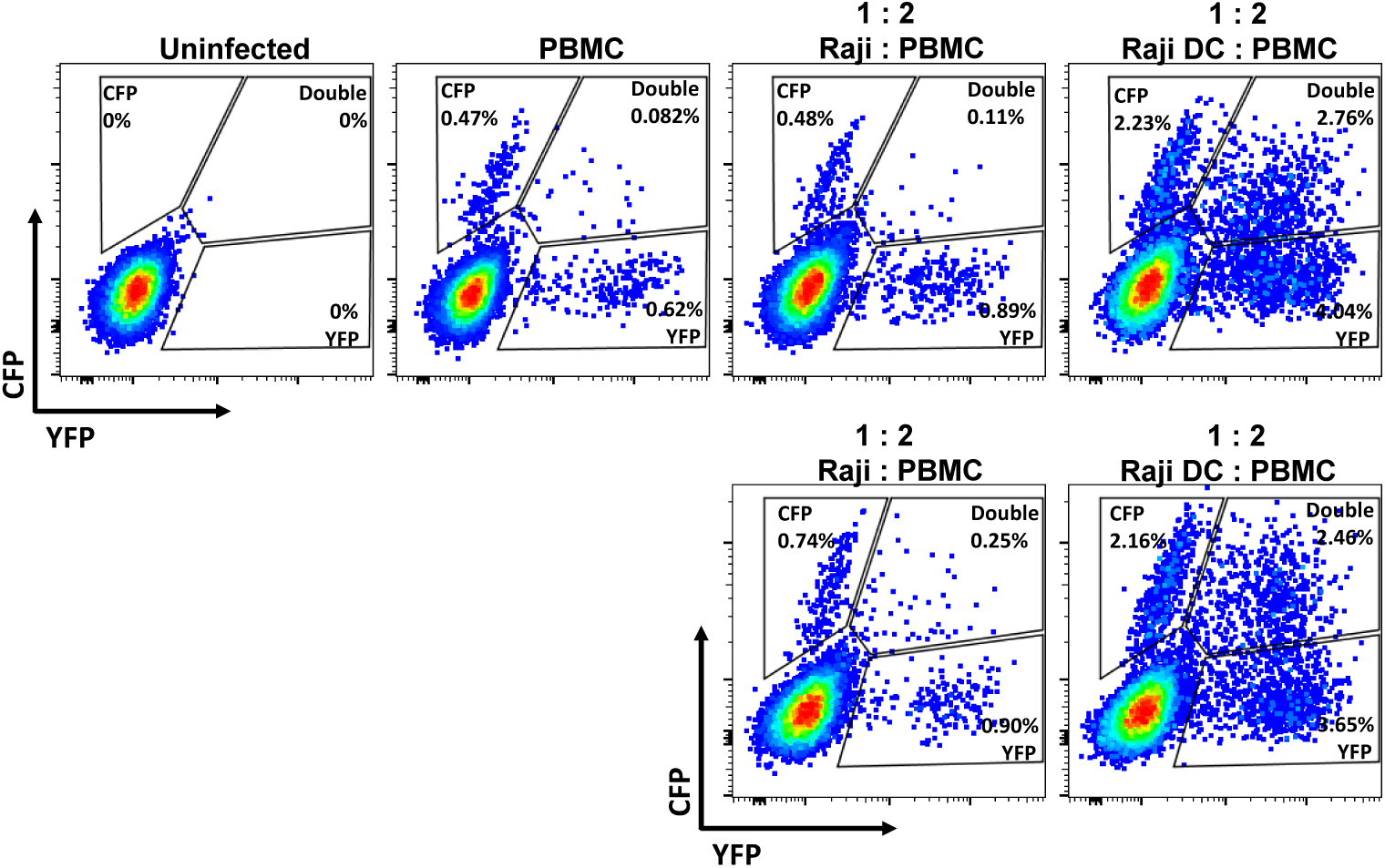
Frequency of double infection in PBMCs is higher that theoretically calculated. Three infections conditions, PBMCs alone, PBMCs + Raji cells and PBMCs + Raji-DC cells, were tested for the level of double infection after incubation with CFP and YFP wildtype producing virus. PBMCs and Raji or Raji-DC cells were combined at a 1:2 ratio. Bottom panel is a repeat with different donor PBMCs. The theoretical probability of double infections, as calculated: [(CFP infection x YFP infection) x 100] < the experimental values.

**Figure S5.**
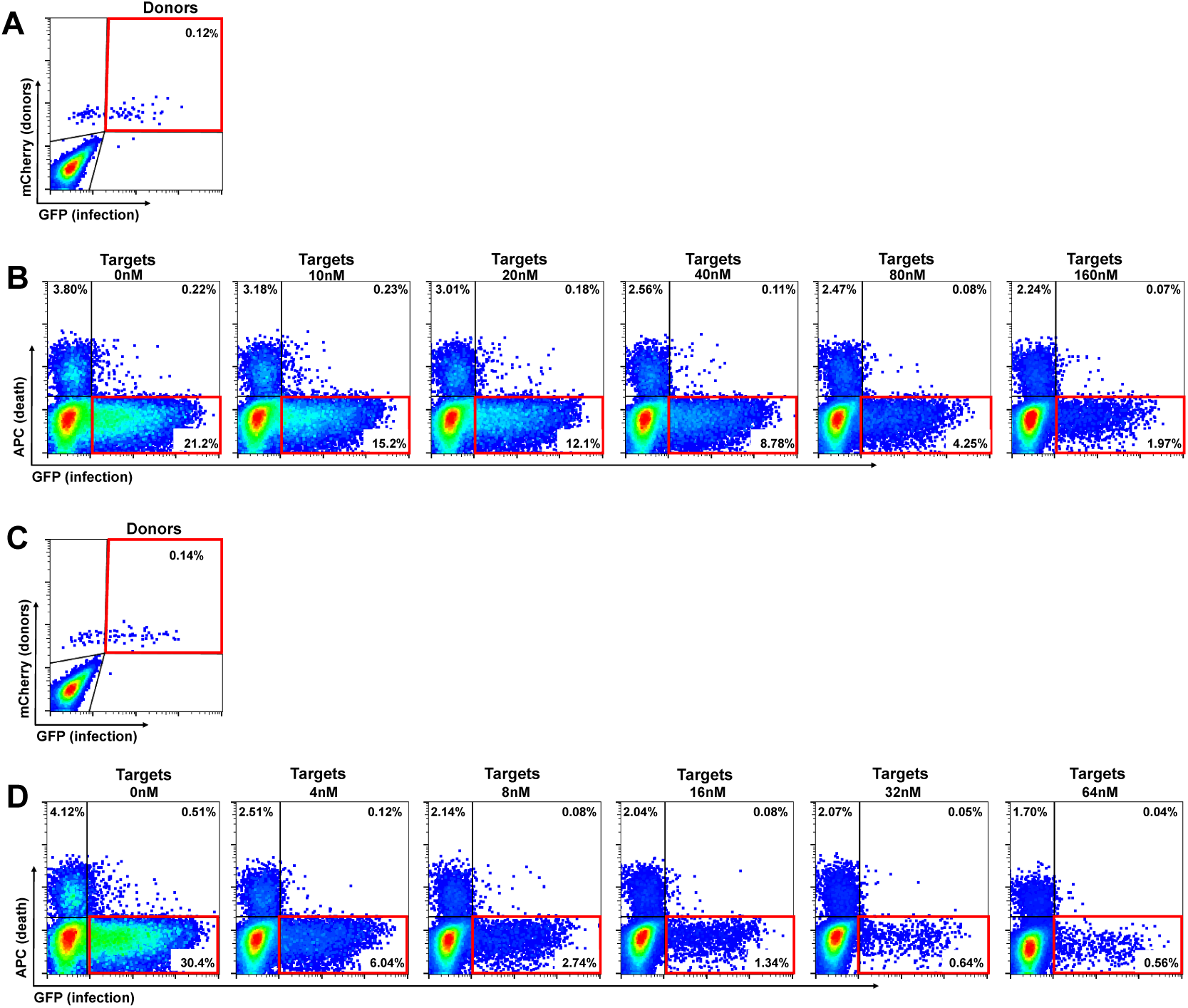
Experimentally measured values of ***R***_*wt*_ and ***R***_*mt*_. (A) The amount of L100I mutant infected G2 donor cells added to E7 target cells as measured straight after donors were added to targets. (B) Target cell infection after 2 days of incubation with L100I mutant infected donors (A) in the presence of different drug concentrations. (C) The amount of wildtype infected G2 donor cells added to E7 target cells as measured straight after donors were added to targets. (D) Target cell wildtype infection after 2 days of incubation with wildtype infected donors (C) in the presence of different drug concentrations. Red squares highlight donors added for (A) and (C) and live infected targets for (B) and (D).

**Table S1.**
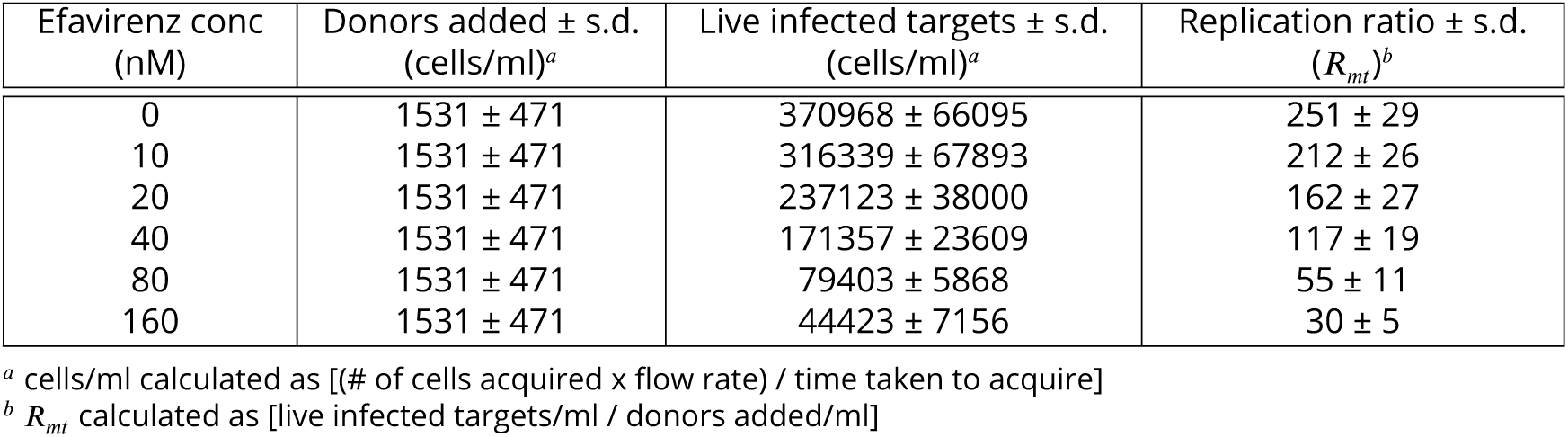
Replication ratio of mutant virus

**Table S2.**
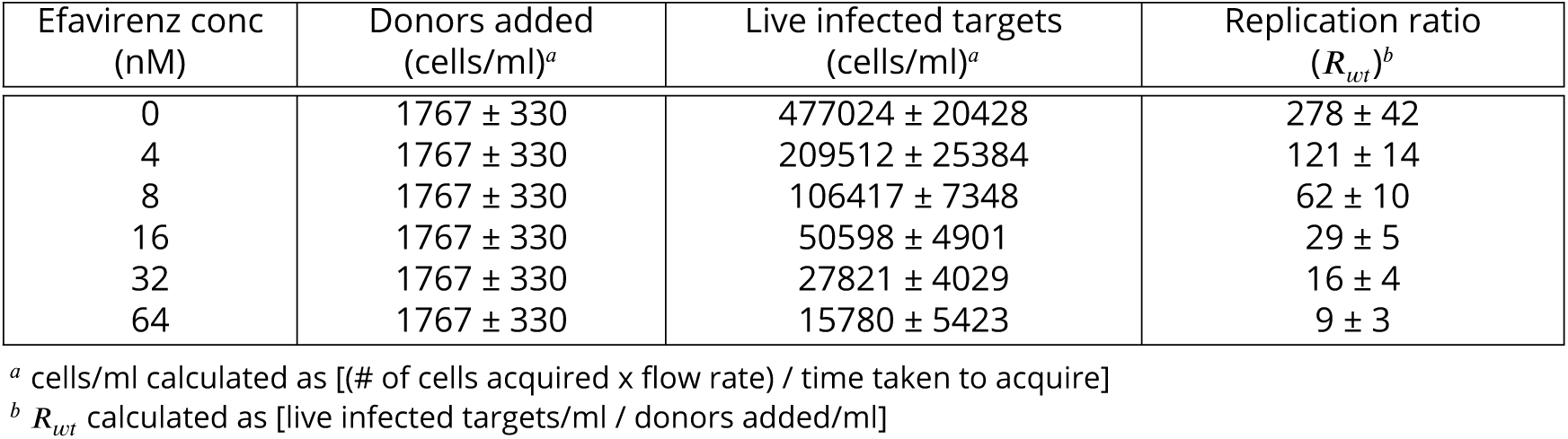
Replication ratio of wildtype virus

